# Genome-wide association study shows developmental robustness control by intestinal maltase via internal environment in *Drosophila*

**DOI:** 10.1101/2025.08.13.669836

**Authors:** Soshiro Kashio, Izumi Enoki, Yuto Yoshinari, Takashi Nishimura, Masayuki Miura

**Affiliations:** Department of Genetics, Graduate School of Pharmaceutical Sciences, The University of Tokyo, 7-3-1 Hongo, Bunkyo-ku, Tokyo 113-0033, Japan; Department of Integrative Bioanalytics, Institute of Development, Aging and Cancer (IDAC), Tohoku University, Sendai 980-8575, Japan; Institute for Molecular and Cellular Regulation, Gunma University, 3-39-15 Showa-machi, Maebashi 371-8512, Japan; Laboratory for Cell Vigor Regulation, National Institute for Basic Biology, Nishigonaka 38, Okazaki, Aichi 444-8585, Japan

## Abstract

Organisms encounter disturbances during development because of genetic variations, environmental shifts, and stochastic noise. However, developmental robustness and canalization buffer these fluctuations to maintain normal development. Nevertheless, understanding the underlying mechanisms that govern this robustness is challenging because of the complex interactions between these factors. In *Drosophila*, the number of scutellar sensory organs (SSO; macrochaetes) derived from the sensory organ precursors (SOPs) in the larval wing disc has been used as a model to study developmental robustness. Although the number of SOPs is strictly regulated by a network of signaling and transcription factors in the imaginal disc, non-intrinsic broader factors such as temperature and energy metabolism additionally influence SSO-number fluctuation. Moreover, the precise molecular mechanisms regulating the systemic control of bristle number remain unknown. In this study, we identify factors controlling bristle robustness by performing genome-wide association studies (GWAS). We observed significant single-nucleotide polymorphisms (SNPs) in the *Maltase* gene cluster and found that the knockdown of *Maltase* genes affected SSO numbers. Furthermore, Maltase-A1(Mal-A1) in the gut regulated insulin signaling systemically, thereby affecting SSO-number fluctuation. These results suggest that Mal-A1 contributes to robustness by modulating glucose availability and *Drosophila* insulin-like peptide 3 (dilp3) level, which affects the SOPs in a nonautonomous manner. This study presents the molecular basis of nutritional regulation of developmental robustness and highlights Maltase as a key mediator.

## Introduction

Organisms encounter disturbances during development because of genetic variation, environmental fluctuations, or stochastic noise ^1^. Despite these intrinsic and extrinsic perturbations, developmental robustness and canalization allow organisms to buffer their phenotypic variability.

An example of genetic robustness is genetic compensation, which involves the activation of other genes to offset the mutation-induced loss-of-function. For example, zebrafish embryos show increased *tfap2c* expression to prevent defects in neural crest cells after the loss of *tfap2a* ^2^. Similarly, in Rel knockout models, the function of the NFκB homolog Rel as a dorsal organizer in zebrafish embryos is compensated by Rela upregulation ^3^. Additionally, robustness to environmental and stochastic noise is linked to specific genes. In *Drosophila*, miR-7 acts as a buffer against temperature fluctuations ^4^. Similarly, mutations in *Drosophila Tps1*, which encodes a trehalose synthesis enzyme, increased variability (developmental robustness and stability) in wing size, especially under variable nutrition conditions. Thus, trehalose metabolism plays a role in developmental robustness and stability through the regulation of glucose homeostasis ^5^.

The scutellar sensory organ (SSO) number in *Drosophila* is an established model for studying developmental robustness. Typically, four macrochaetes (bristles) are found on the scutellum, but some individuals exhibit five or more macrochaetes (Fig. 1A). These bristles are originated from sensory organ precursors (SOPs) in the wing discs of third-instar larvae through asymmetric cell division ^6,7^. SOP selection within proneural clusters (PNCs) is primarily controlled by the Notch signaling pathway (Fig. 1B) with the expression of the proneural genes *achaete* (Ac) and *scute* (Sc), which encode basic helix–loop–helix (bHLH) transcription factors ^8,9^. These bHLH factors target the transcription factor Senseless (sens), which feeds back Ac and Sc expression, leading to the sensory organ fate ^10^.

**Fig. 1.**
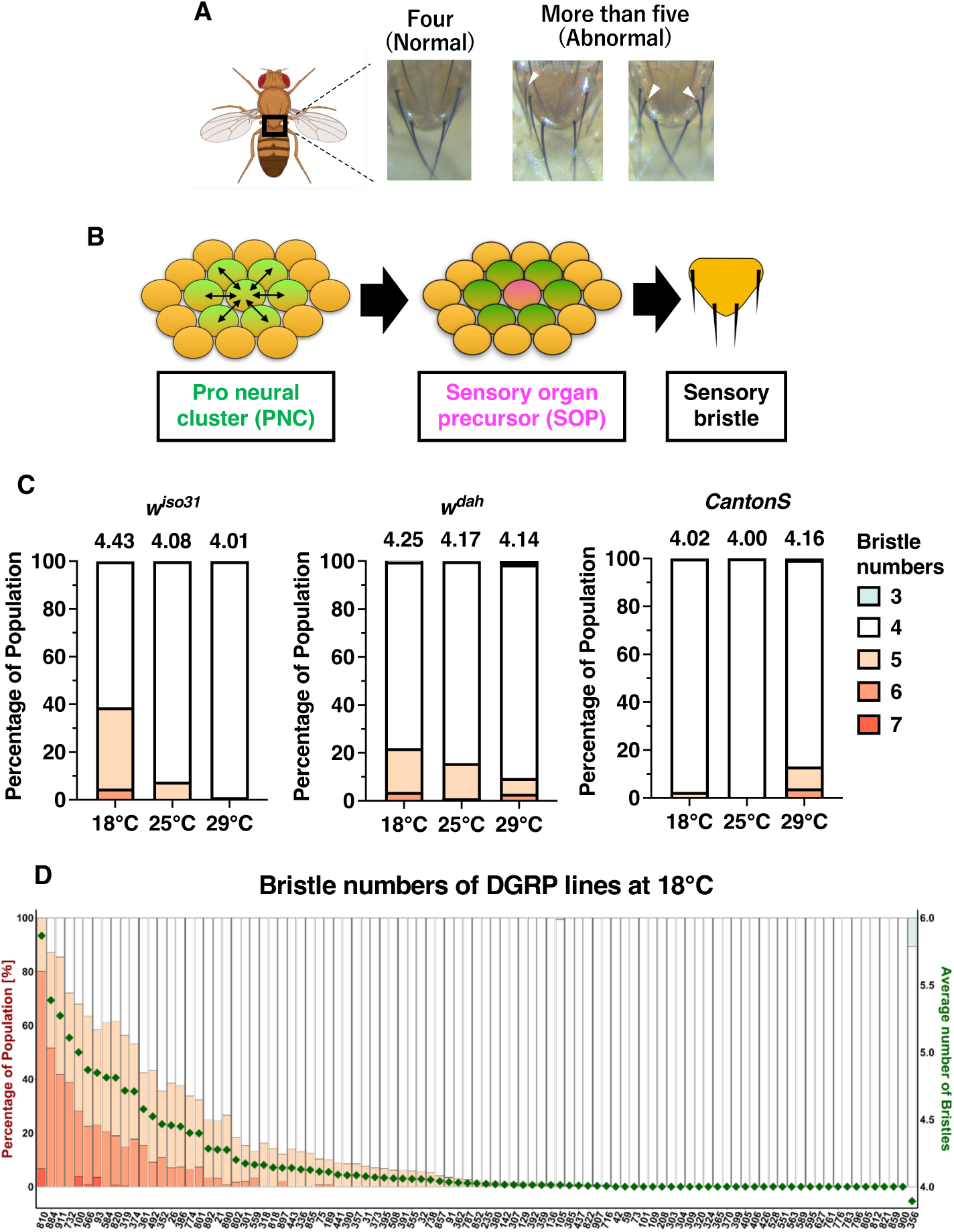
Genome-Wide Association Study (GWAS) of developmental robustness using bristle number as indicator. **(A)** Photograph showing macrochaetes (large bristles) within the scutellum region of *Drosophila*. Although the typical number of bristles is 4, some individuals exhibit abnormal number bristles (white arrowheads) such as ≥ 5. **(B)** In larval *Drosophila*, proneural clusters (PNCs) within the wing imaginal disc are specified into sensory organ precursors (SOPs). **(C)** Bristle numbers in wild-type strains *w^iso31^*, *w^dah^*, and *Canton-S* reared at 18, 25, and 29 °C. The mean bristle number is indicated above each graph. From left to right, n=214, 2773, 95 (*w^iso31^*), 429, 198, 138 (*w^dah^*), 201, 120,152 (*Canton-S*). **(D)** Proportion and mean bristle number for each DGRP line. The number of bristle in the females were counted at 18 °C in 95 DGRP lines. The x-axis represents the DGRP lines, left y-axis indicates the bristle number populations, and right y-axis represents the mean bristle number (green dots).

The precise number of SSOs critically depends on the level of proneural gene expression, ^11^ which is regulated by factors within the PNC. For example, the serine/threonine kinase Shaggy (Sgg) is a GSK-3β ortholog that negatively regulates Sc through phosphorylation ^12^. Wingless (Wg) is a WNT ortholog that inhibits Sgg to support SOP development ^13^. Caspases play a non-apoptotic role to aid in SOP formation by activating the Sgg isoform Sgg46 ^14^. Additionally, miR-9a reduces *sens* expression to prevent the formation of extra bristles ^15–17^. In addition to these pathways, epigenetic factors are also involved, such as the in the promotion of SOP formation by the histone methyltransferase SETDB1 ^18^.

In addition to the intrinsic factors that are specifically required for sensory organ production, metabolic and translational regulators also affect the SSO numbers. For instance, the genetic removal of insulin-producing cells (IPCs) or mutations in the ribosomal protein *RpS* reduced the increased bristle numbers in *miR-9a* mutants, suggesting that slower metabolism and translation reduce developmental errors ^19^. Although environmental factors such as temperature and nutrition have been shown to influence SOP generation ^16,20^, the underlying molecular mechanisms have not been elucidated.

In this study, we performed genome-wide association studies (GWAS) to identify genes linked to significant single-nucleotide polymorphisms (SNPs) associated with bristle number, followed by RNAi screening. This approach showed that the maltase gene cluster—specifically Maltase-A1 (Mal-A1) in the gut— is a regulator that influences SSO robustness through insulin signaling. These findings uncover a previously unrecognized link between digestive enzymes and developmental robustness, providing new insight into how systemic metabolic cues shape SSO robustness.

## Results

### Bristle numbers varied in wild-type lines based on rearing temperature

We investigated the impact of genetic background and environment on SSO (represented by macrocheate or bristle) numbers by first examining three wild-type lines at rearing temperatures of 18, 25, and 29 °C. We found that the effect of temperature on bristle number varied between the wild-type lines (Fig. 1C). As ectopic bristle position shifts with temperature^21^, we analyzed the bristle location in detail and categorized the ectopic bristles as Anterior (near the anterior scutella, aSC), Posterior (beneath the posterior scutella, pSC), or Interval (between the aSC and pSC) (Supplementary Figure 1A). As Interval bristles were rare, we combined them with the Anterior bristles for analysis (Supplementary Figure 1B). In concurrence with the findings of a previous study ^21^, Anterior or Interval bristles and no posterior bristles were observed in the analyzed lines at 18 and 25 °C, whereas a high number of Posterior bristles were observed at 29 °C (Supplementary Figure 1B and 1C). Notably, the ectopic Posterior bristles were mostly observed at 29 °C in the *CantonS* line, whereas the Anterior or Interval ectopic bristles were most observed at 18 or 25 °C in the *w^iso31^*and *w^dah^* lines (Supplementary Figure 1C). These findings suggest that the ectopic Posterior bristles observed at 29 °C are formed as a result of underlying mechanisms that are distinct from those governing the ectopic Anterior and Interval bristles observed at 18 and 25 °C. Moreover, rearing at 29 °C reduced the eclosion rates, suggesting that the ectopic Posterior bristles appear at 29 °C because of stress from heat. Furthermore, this distinguishes the bristles formed under high-temperature stress from those formed under relatively low temperatures. Further analysis was performed using the phenotype at 18 °C as an indicator because a greater increase in bristle was observed at 18 °C than at other temperatures.

### GWAS using DGRP lines identified SNPs associated with bristle numbers

We identified the genetic factors controlling bristle numbers by performing a GWAS using the *Drosophila* Genetic Reference Panel (DGRP), which contains approximately 200 wild-type lines ^22,23^. We compared the traits and genomes across these lines and statistically identified the SNPs associated with the traits. Furthermore, we identify the genetic loci associated with bristle number variability using this approach.

We examined 95 lines at 18 °C. Some lines showed a high proportion of individuals with more than five bristles, indicating a substantial genetic influence on bristle number (Fig. 1D).

The dorsocentral (DC) bristles are structures that occur on the notum and acts as a sensory organ. They typically exhibit a consistent count of four bristles and comprise the anterior DC (aDC) and posterior DC (pDC) bristles (Supplementary Figure 2A). We compared the DC and SC bristle-number regulation across the DGRP lines. We found that although both SC and DC bristle numbers tended to increase, the correlation between the DC- and SC-bristle numbers was weak between the different lines, with R^2^ = 0.07 at 18 °C (Supplementary Figure 2B). This suggests that although temperature affected both bristle types, the regulation of bristle numbers varied for the DC and SC bristles because of the genetic background.

We conducted a GWAS based on the average SC bristle numbers at 18 °C in the DGRP lines to explore the genetic factors that contribute to SC bristle-number robustness and identified several SNPs associated with SC-bristle number (Fig. 2A). The Q–Q plots confirmed that the distribution of the SNPs matched the expected probability distribution (Supplementary Figure 2C).

**Fig. 2.**
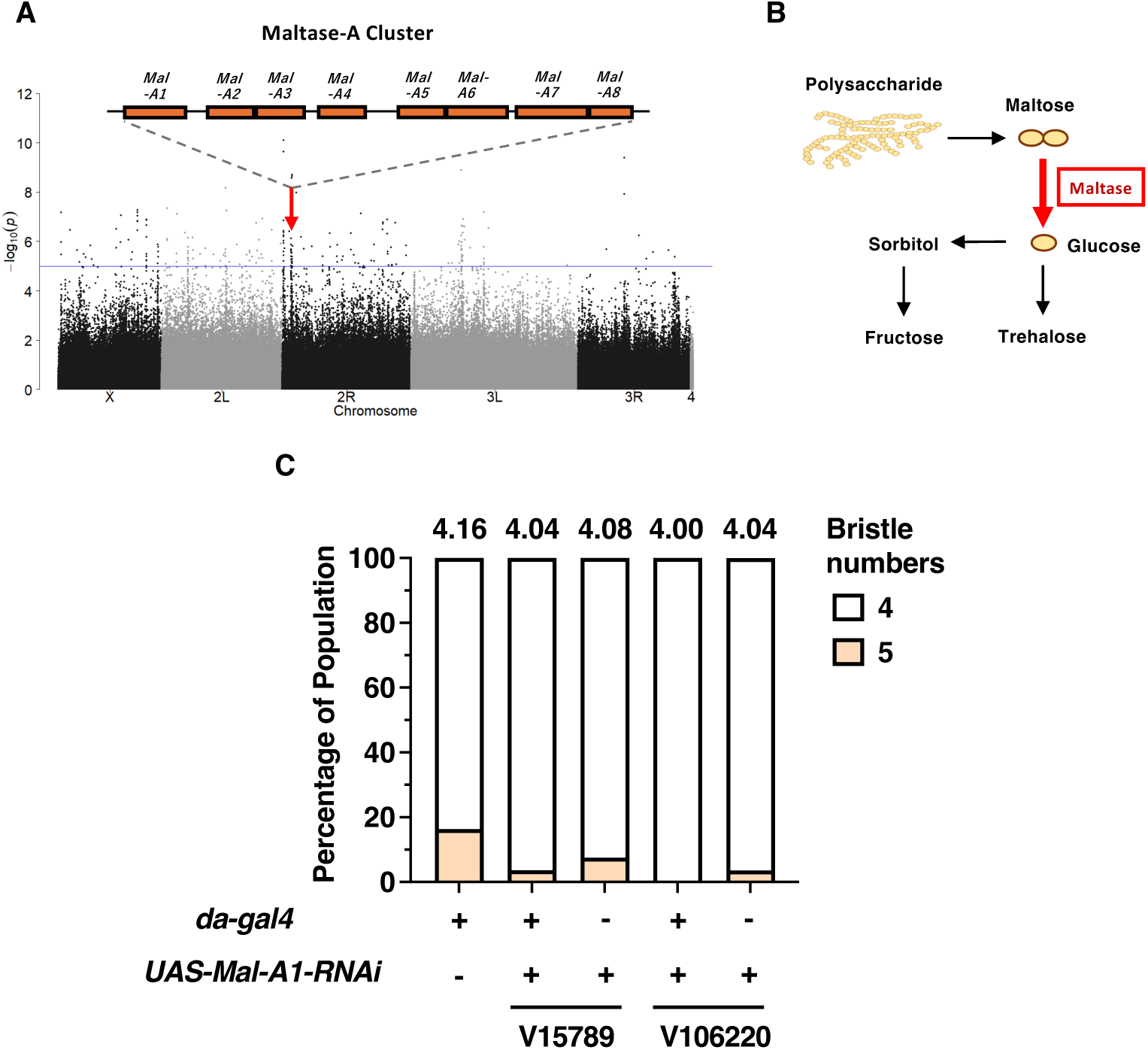
Reduction of bristle numbers following whole-body knockdown of *Mal-A1*. **(A)**Manhattan plots showing GWAS results for mean bristle numbers at 18°C. The blue line represents -log10(p) = 5, and the red line represents the significance threshold calculated using Bonferroni correction and corresponding to a p-value of 0.05 divided by the number of SNPs analyzed for each condition (p = 3.33e-08). The genomic location of the maltase cluster in the GWAS Manhattan plot. The mean bristle number is indicated above each graph. **(B)** Schematic illustration of maltase function in sugar metabolism. **(C)** Reduction in bristle numbers following *Mal-A1* knockdown in the whole body at 25 °C. Two different *Mal-A1* knockdown lines (V15789 and V106220) are examined. The mean bristle number is indicated above each graph. From left to right, n=55, 111, 40, 37, 56.

### RNAi screening identified Mal-A1 as a systemic key contributor to bristle number

We verified the hypothesis that genes harboring the SNPs linked to bristle number contribute to bristle formation by analyzing the genes that showed moderate or higher expression levels in the third larval stage (based on FlyBase modENCODE Development RNA-Seq), followed by an RNAi screening of 43 candidate genes at 18 °C (Table 1). In the first screening, progeny from *da>RNAi* crosses were compared with both parental *UAS-RNAi* and *da-gal4* lines, and lines showing altered SC bristle numbers were selected. These candidates were re-tested in the second screening using the same comparison to confirm reproducibility. In the third screening, the selected lines were tested again using different sets of parental controls to exclude background effects. Through this stepwise screening using the whole-body *da-gal4* driver, we identified *Maltase-A1* (*Mal-A1*) as a contributor to bristle number in the third screening (Table 1).

**Table 1.** RNAi screening of candidate genes from GWAS of whole-body knockdown at 18 °C. Results of whole-body RNAi screening of candidate genes identified at 18 °C in the GWAS. The p-values shown in the table represent the smallest p-value among the SNPs within each gene. *Da-gal4* was used for whole-body induction. The table shows sample sizes (n), mean bristle numbers, and the increase or decrease in bristle numbers following gene knockdown compared with those of both controls: parental UAS lines and parental gal4 line. For the 1^st^ screening, the bristle number of progeny *da>RNAi* lines were compared with that of the corresponding parent *UAS-RNAi* lines (without driver) and *da-Gal4* line (without UAS). The lines with altered bristle numbers were selected (average number of SC bristles in *da-Gal4* line was 4.16 (n=55)) For the 2^nd^ screening, the results were confirmed by repeating the same comparison with the lines selected during the first screening. For the 3^rd^ screening, the bristle number of the progeny *da>RNAi* lines were compared with those of different sets of parent *UAS-RNAi* lines (without driver) and *da-Gal4* line (without UAS). Once again, the lines with the altered bristle numbers were selected. The selected *UAS-RNAi* lines are indicated in blue, and parental *da-Gal4* line is marked in green.

### Knockdown of multiple *maltase* genes suppressed the increase in bristle number

*Mal-A1* encodes a maltase enzyme that breaks down maltose into glucose (Fig. 2B). Knockdown of *Mal-A1* suppressed the increase in bristle number (Fig.2C). The *Drosophila* genome contains ten maltase genes, with *Mal-A1-8* clustered on chromosome 2R (Fig. 2A). GWAS analysis of other maltase genes showed several SNPs that were linked to bristle number in the *maltase* gene cluster, including variants located in coding exons, introns, and untranslated regions (UTRs), with several clustered in coding regions that could potentially affect protein function (Table 2). Various types of SNPs were detected in the Maltase-A cluster, and several DGRP lines with extra bristles showed alternative alleles in the Maltase-A genes (Table 2). We investigated their contributions by performing whole-body knockdown of other maltase genes, namely, *Mal-A3* and *Mal-A4* (Supplementary Figure 3A). Knockdown efficiency was confirmed for *Mal-A1*, *Mal-A3*, and *Mal-A4* using *da-Gal4* (Supplementary Figure 3B).

**Table 2.** SNPs in the Maltase cluster associated with bristle number in DGRP lines. The average number of macrochaetae at 18 °C in each DGRP line and a list of SNPs in the Maltase gene cluster that exhibited strong associations with macrochaete number. The locus of SNPs, p value in manhattan plots, and sequence ontology are described. The alleles are indicated using ‘0’ for reference (green), ‘–’ for missing (green), and ‘2’ for alternative (yellow).

### Whole-body *Mal-A1* knockdown caused metabolic alterations

Maltases are highly expressed in the gut, where they facilitate glucose production through maltose digestion; hence, we hypothesized that Mal-A1 contributes to bristle number robustness by systemically regulating the glucose levels. We investigated the metabolic changes induced by whole-body *Mal-A1* knockdown using liquid chromatography–mass spectrometry (LC–MS), gas chromatography–mass spectrometry (GC–MS), and principal component analysis, which showed a distinct separation between *Mal-A1* knockdown and control samples (Fig. 3A). Volcano plot analysis further confirmed the significant differences (Supplementary Figure 4), particularly in sugar metabolism. We found that maltose level increased, whereas that of glucose decreased (Fig. 3C). In addition to maltose-related sugars, the levels of arabitol, ribitol, and ribose increased and those of isomaltose and sophorose decreased. This indicates that Mal-A1 knockdown affects a wide range of sugars (Supplementary Figure 4).

**Fig. 3.**
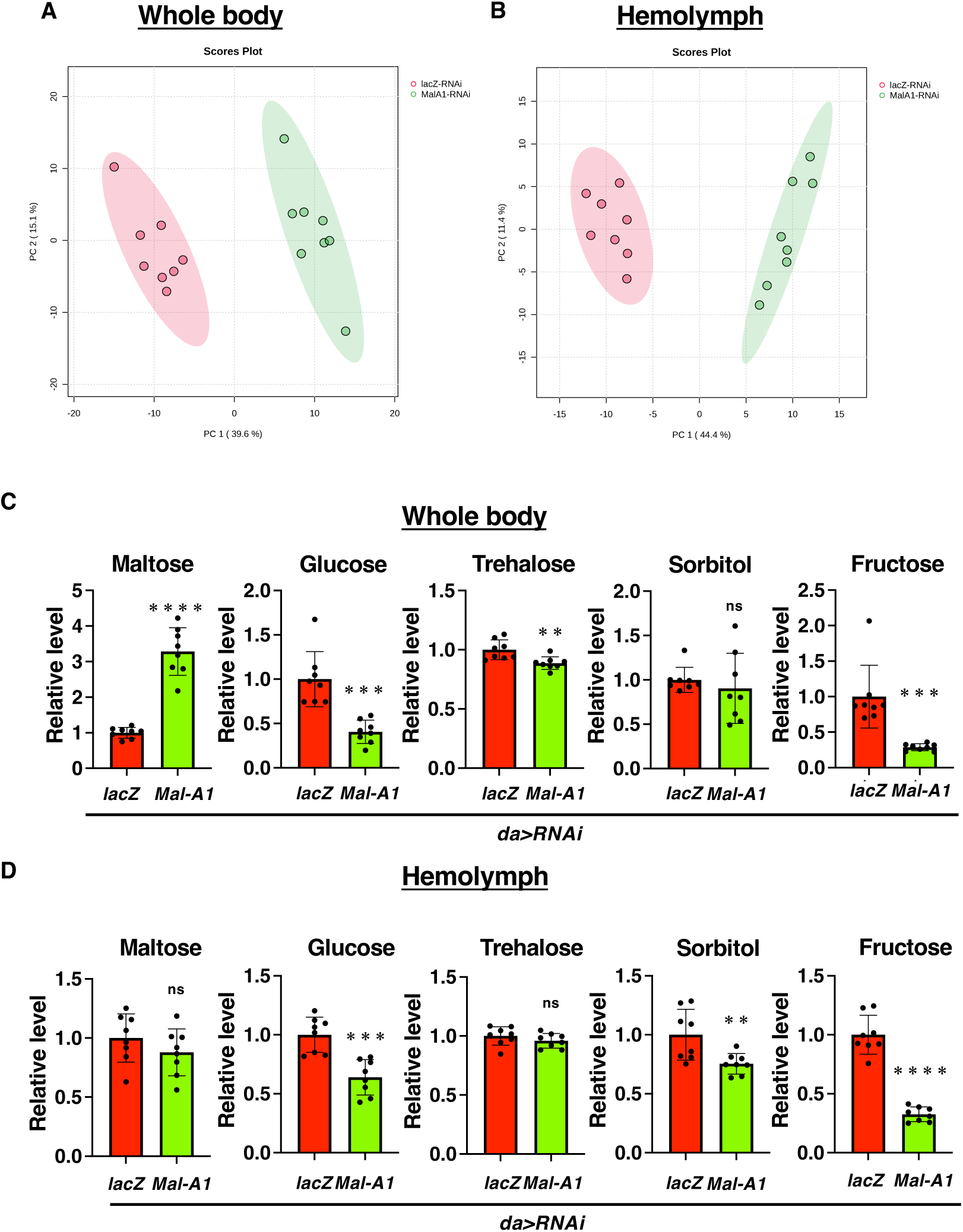
Metabolic changes induced by whole-body knockdown of *Mal-A1*. **(A, B)** Principal component analysis of water-soluble metabolites measured using LC-MS following *Mal-A1* knockdown (V15789 line): (**A**) whole-body samples and (**B**) hemolymph samples. **(C, D)** Changes in sugar metabolite levels associated with maltose and glucose metabolism measured using LC-MS and GC-MS following *Mal-A1* knockdown: (**C**) whole-body samples and (**D**) hemolymph samples. The quantity of each sugar is represented as peak area of mass spectrometric metabolome analysis.

Additionally, we quantified the metabolites in the hemolymph (Fig. 3B, 3D, and Supplementary Figure 5). Unlike that observed in the whole-body samples, maltose level did not change, which suggests that the increase in maltose was likely due to its accumulation within specific tissues such as the intestine. However, similar to that in the whole-body samples, glucose level decreased in the hemolymph (Fig. 3D). The decrease in glucose level in the hemolymph aligns with the function of Mal-A1 and suggests broad effects on sugar metabolism.

### *Mal-A1* knockdown reduced *dilp3* expression in the CNS

Next, we investigated the manner in which reduction in blood sugar levels in the hemolymph affected the bristle number. Considering that blood glucose levels influence insulin production, we investigated whether Mal-A1 influences insulin-producing cells (IPCs) in the larval central nervous system (CNS). We assessed *dilp2, dilp3,* and *dilp5* expression in the CNS following *Mal-A1* knockdown. Although *dilp2* and *dilp5* levels remained unchanged, *dilp3* expression decreased in the CNS (Fig. 4A). Furthermore, brain immunostaining showed decreased Dilp3 signal after *Mal-A1* knockdown and no change in Dilp2 signal (Fig. 4B).

**Fig. 4.**
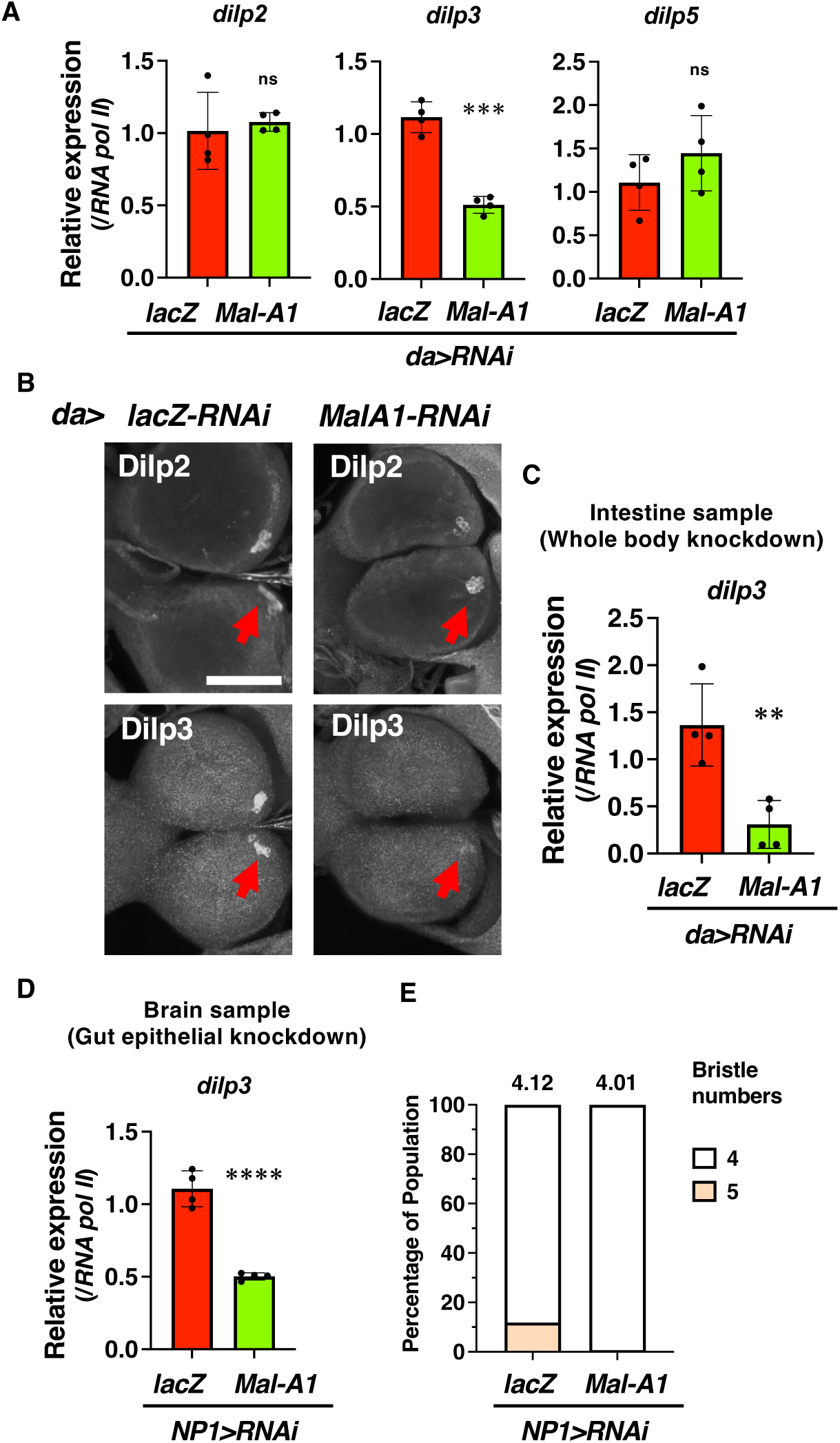
Changes in gene expression of CNS *dilps* and developmental robustness reflected in bristles via gut *Mal-A1* knockdown. **(A)** Comparison of *dilp2*, *dilp3*, and *dilp5* expression in the CNS at 96 h after egg laying (AEL) at 25 °C after whole-body knockdown of *Mal-A1* (V15789 line). **(B)** Immunostaining of Dilp2 and Dilp3 in the CNS after whole-body *Mal-A1* knockdown; arrows indicate IPCs. Dilp3 signal decreased after *Mal-A1* knockdown. Scale bar, 75 µm. **(C)** Reduced *dilp3* expression in the intestine after whole-body *Mal-A1* knockdown. **(D)** Reduced *dilp3* gene expression in the brain after *Mal-A1* knockdown in gut epithelial cells. **(E)** Reduced bristle numbers following *Mal-A1* knockdown in gut epithelial cells.

Additionally, as *dilp3* is also expressed in the intestine, we examined its gene expression in the intestine after *Mal-A1* knockdown. Intestinal *dilp3* expression decreased following whole-body *Mal-A1* knockdown (Fig. 4C), indicating that intestinal *dilp3* expression is also influenced by Mal-A1. Additionally, enterocyte-specific *Mal-A1* knockdown reduced *dilp3* expression in the CNS (Fig. 4D). This result supports the significance of Mal-A1 and gut glucose breakdown in regulating Dilp3 expression and possibly bristle-number robustness. Similar to that observed after whole-body knockdown, enterocyte-specific *Mal-A1* knockdown increased the proportion of flies with four bristles (Fig. 4E).

### Increase in cold-induced insulin level was consistent with increase in bristle number

InR regulates the expression of the proneural genes *achaete* and *scute* ^24^. *InR* overexpression in PNC increased the bristle number, whereas *InR* knockdown reduced the bristle number ^24^. The findings of the current study concur with these results, which suggest that maltose breakdown and control of glucose levels in the hemolymph by *Mal-A1* in the intestine influence bristle number via Dilp3-mediated insulin signaling in SOP cells. As systemic insulin signaling is apparently crucial in regulating bristle number, we investigated the correlation between bristle number and Dilps under physiological conditions that alter the expression of Dilps. Specifically, we compared *dilp* expression levels under different temperature conditions (25 and 18 °C) that produced a change in the number of bristles in the wild-type *w^iso31^* flies (Fig. 1C). At the late third-instar larval wandering stage, *Drosophila* larvae leave their food source in search of a pupation site. At 25 °C rearing temperature, they reach this stage at approximately 96 h post-laying ^25^. The equivalent wandering stage for 18 °C occurs at approximately 216 h post-laying. We compared *dilp* mRNA expression in the CNS of wild-type *w^iso31^* larvae at these stages—96 h post egg laying at 25 °C and 216 h post-laying at 18 °C. The results showed that both *dilp2* expression and bristle number were higher at 18 °C than at 25 °C. Similarly, *dilp3* showed higher expression at 18 °C; however, *dilp5* did not (Fig. 5A and 1C). This trend is confirmed by previously reported results of increased IPC activity and Dilp secretion 6 h after cold exposure ^26^. These results suggest a consistent relationship between the decrease in extra-bristles observed as a result of reduced *dilp3* expression due to *Mal-A1* knockdown and the increase in extra-bristles observed as a result of elevated *dilp2* and *dilp3* expression at lower temperatures (Fig. 5B).

**Fig. 5.**
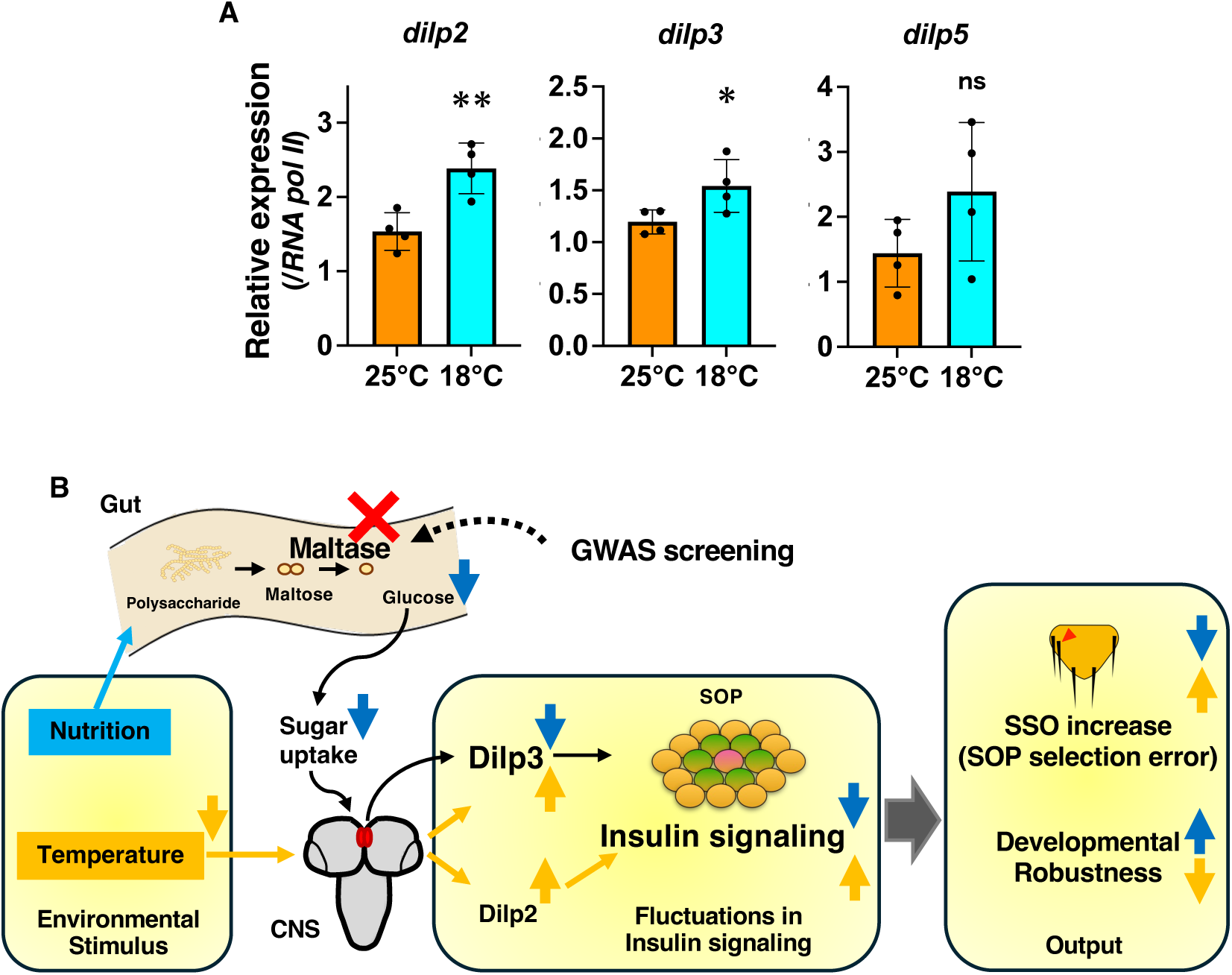
Changes in gene expression of CNS *dilps* and developmental robustness reflected in bristles via sugar metabolism during temperature variations. (**A**) Comparison of CNS expression of *dilp2*, *dilp3*, and *dilp5* in *w^iso31^* at 96 h AFT at 25 °C or at 216 h AFT at 18 °C. **(B)** Model of developmental robustness mediated by insulin signaling induced by maltase and temperature. Maltases identified in the GWAS regulate sugar metabolism in the gut, influencing *dilp3* expression in the central brain, thereby controlling developmental robustness as reflected in bristle numbers. Additionally, temperature-dependent changes in *dilp2* and *dilp3* expression affected bristle numbers.

## Discussion

This study has identified maltase enzymes—particularly Mal-A1—as key regulators of SSO (macrochaete or bristle)-number robustness in *Drosophila*, acting primarily through the modulation of insulin signaling. Notably, *Mal-A1* was expressed in the intestine, and it modulated insulin signaling and glucose metabolism. This finding highlights its importance in nutrient-dependent developmental stability (Fig. 5B).

The expression of maltase along with other digestive enzymes such as amylase is regulated by feedback from the metabolic state of the fat body, which is the *Drosophila* organ that is analogous to the mammalian liver and adipose tissue ^27^. This feedback suggests that maltase plays a role in modulating glucose metabolism and insulin signaling by adjusting the intestinal glucose supply in response to nutritional conditions. The data consistently indicate that Mal-A1 influences glucose availability, thereby modulating insulin production and signaling. Additionally, we found that *Mal-A1* knockdown specifically led to reduced *dilp3* levels in the IPCs and gut and enhanced bristle-number robustness (Fig. 4). As *dilp3* expression is sensitive to circulating glucose ^28^, glucose regulation by Mal-A1 highlights its role in maintaining robust developmental outcomes via insulin signaling.

In this study, the knockdown of a single maltase reduced *dilp3* expression and suppressed the increase in bristle number (Fig. 2C and 4E). However, *Mal-A1* knockdown notably altered sugar metabolism (Fig. 3). Generally, the genes in clusters are expected to exhibit compensatory mechanisms. The lack of compensation observed in this study may indicate the presence of a maltose threshold that determines systemic glucose levels, implying a vulnerability to developmental robustness.

The importance of glucose metabolism in developmental robustness has been explored by analyzing the relationship between trehalose and wing-size robustness under nutritional fluctuations, and trehalose stabilized glucose concentrations, thereby reducing inter-individual variation in wing size ^5^. In contrast, *Mal-A1* knockdown in this study via the whole-body Gal4 driver (*da-gal4*) resulted in only a minor reduction in trehalose, suggesting that fluctuation in glucose levels exert a direct effect on bristle number via insulin signaling. However, the effect of maltase on inner-individual traits such as bilateral symmetry in wing size remains unknown when using bristle number as the inter-individual indicator and warrants further investigation.

Furthermore, these results highlight the role of insulin signaling in developmental robustness under environmental stress, particularly that of low blood glucose levels caused by *Mal-A1* knockdown or low temperature (Fig. 5B). Although low temperatures may induce metabolic slowdown and subsequent developmental stability, low temperatures elevated the expression of insulin-like peptides encoded by *dilp2*, *dilp3*, and *dilp5* ^29^, probably to sustain metabolic activity. During SSO development, increased insulin signaling increased proneural gene expression via the InR/FOXO pathway ^24^. Although TOR functions downstream of insulin signaling to regulate translation, the TOR pathway does not affect bristle number ^24^. This suggests that the increase in SSO number and compromised developmental robustness under low-temperature conditions are likely caused by the upregulation of Dilps rather than by a general metabolic slowdown or reduced translation.

Under IPC ablation conditions, which reduce insulin production, the increase in bristle number seen in *miR-9a* mutants was suppressed, and the proportion of individuals with a normal number of SSOs increased ^19^. This observation coincides with the reduction in bristle number in *Mal-A1-RNAi* flies as a result of Dilp3 downregulation or with the increase in bristle number at low temperatures as a result of Dilp2 and Dilp3 upregulation (Fig. 2C and 1C). Thus, this study presents a concrete mechanism by which internal physiological states influence developmental robustness. Additionally, it highlights the systemic role of maltase upstream of insulin signaling.

Future studies should investigate whether maltase-mediated glucose metabolism affects other developmental stability markers such as bilateral symmetry to provide insights into the physiological adaptations that sustain trait stability across fluctuating environmental conditions. Additionally, the integration of maltase with insulin signaling needs to be further investigated to elucidate the broader regulatory networks that are vital for developmental robustness. This study has established a framework for exploring the manner in which intestinal maltase-driven glucose metabolism interacts with hormonal signaling to support robustness across diverse conditions.

## Materials and Methods

### Fly strains and rearing conditions

Unless specified otherwise, the flies used in this study were maintained in an incubator at 25 °C. Sorting and crosses were performed at room temperature. The following fly strains were used:

**(1) Wild-type strains used for phenotypic analysis:**

- w^iso31^
- w^dah^
- CantonS

**(2) Genome-wide association study (GWAS):**

• *Drosophila Genetic Reference Panel (DGRP)*

GWAS was performed at 18 °C using the following strains:

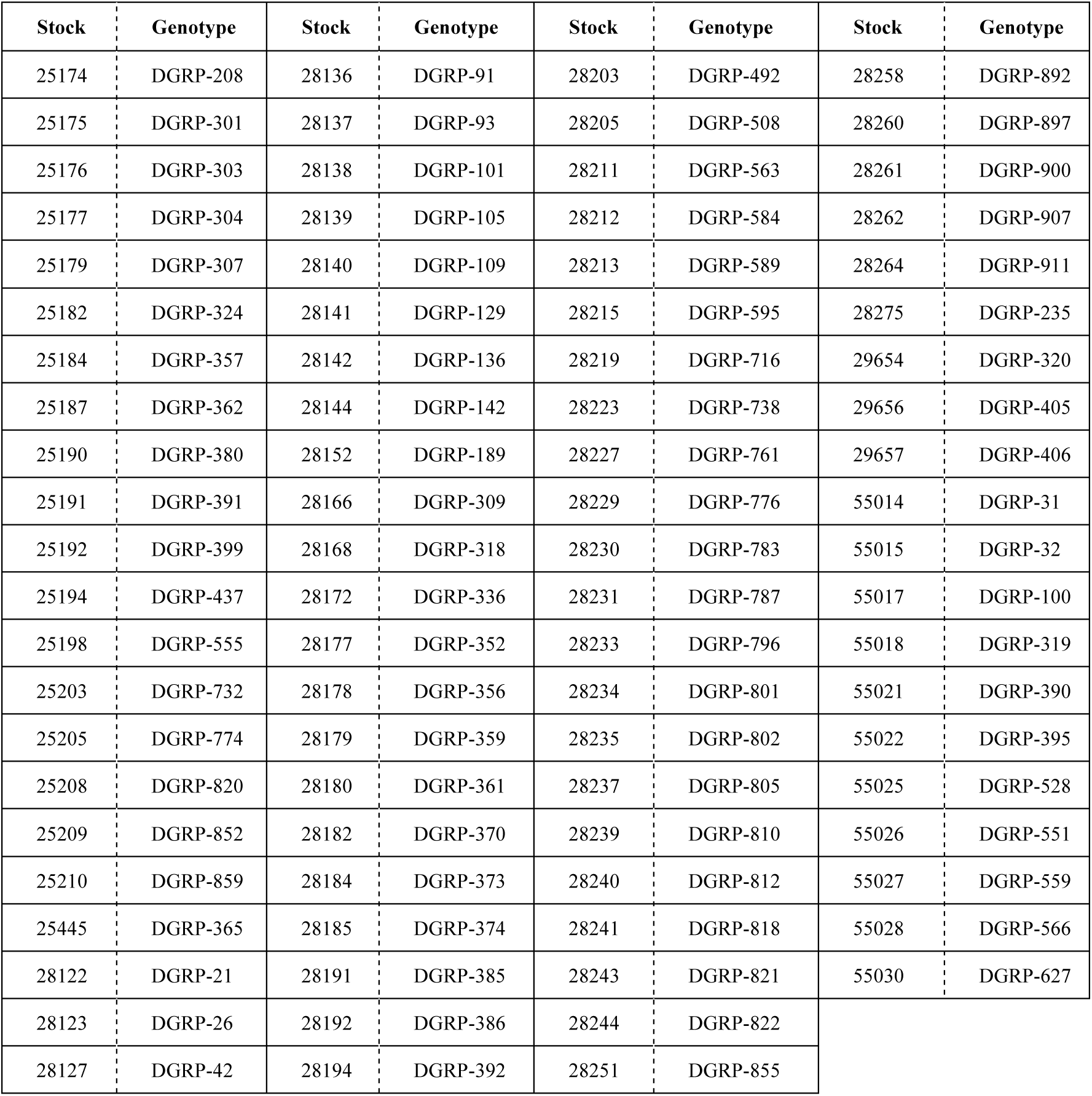
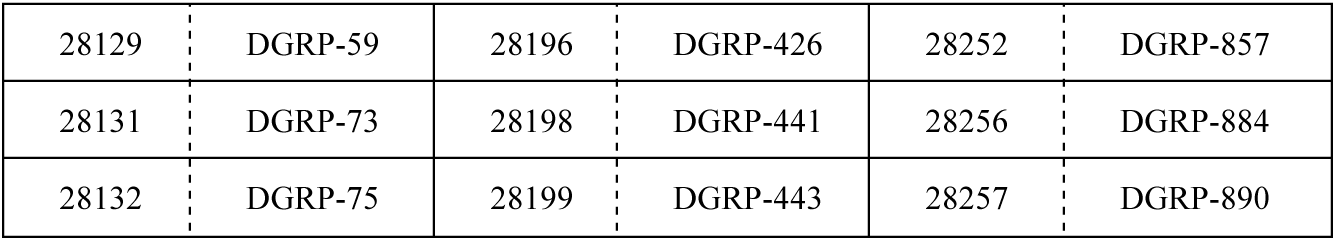

| Stock | Genotype | Stock | Genotype | Stock | Genotype | Stock | Genotype |
| --- | --- | --- | --- | --- | --- | --- | --- |
| 25174 | DGRP-208 | 28136 | DGRP-91 | 28203 | DGRP-492 | 28258 | DGRP-892 |
| 25175 | DGRP-301 | 28137 | DGRP-93 | 28205 | DGRP-508 | 28260 | DGRP-897 |
| 25176 | DGRP-303 | 28138 | DGRP-101 | 28211 | DGRP-563 | 28261 | DGRP-900 |
| 25177 | DGRP-304 | 28139 | DGRP-105 | 28212 | DGRP-584 | 28262 | DGRP-907 |
| 25179 | DGRP-307 | 28140 | DGRP-109 | 28213 | DGRP-589 | 28264 | DGRP-911 |
| 25182 | DGRP-324 | 28141 | DGRP-129 | 28215 | DGRP-595 | 28275 | DGRP-235 |
| 25184 | DGRP-357 | 28142 | DGRP-136 | 28219 | DGRP-716 | 29654 | DGRP-320 |
| 25187 | DGRP-362 | 28144 | DGRP-142 | 28223 | DGRP-738 | 29656 | DGRP-405 |
| 25190 | DGRP-380 | 28152 | DGRP-189 | 28227 | DGRP-761 | 29657 | DGRP-406 |
| 25191 | DGRP-391 | 28166 | DGRP-309 | 28229 | DGRP-776 | 55014 | DGRP-31 |
| 25192 | DGRP-399 | 28168 | DGRP-318 | 28230 | DGRP-783 | 55015 | DGRP-32 |
| 25194 | DGRP-437 | 28172 | DGRP-336 | 28231 | DGRP-787 | 55017 | DGRP-100 |
| 25198 | DGRP-555 | 28177 | DGRP-352 | 28233 | DGRP-796 | 55018 | DGRP-319 |
| 25203 | DGRP-732 | 28178 | DGRP-356 | 28234 | DGRP-801 | 55021 | DGRP-390 |
| 25205 | DGRP-774 | 28179 | DGRP-359 | 28235 | DGRP-802 | 55022 | DGRP-395 |
| 25208 | DGRP-820 | 28180 | DGRP-361 | 28237 | DGRP-805 | 55025 | DGRP-528 |
| 25209 | DGRP-852 | 28182 | DGRP-370 | 28239 | DGRP-810 | 55026 | DGRP-551 |
| 25210 | DGRP-859 | 28184 | DGRP-373 | 28240 | DGRP-812 | 55027 | DGRP-559 |
| 25445 | DGRP-365 | 28185 | DGRP-374 | 28241 | DGRP-818 | 55028 | DGRP-566 |
| 28122 | DGRP-21 | 28191 | DGRP-385 | 28243 | DGRP-821 | 55030 | DGRP-627 |
| 28123 | DGRP-26 | 28192 | DGRP-386 | 28244 | DGRP-822 |  |  |
| 28127 | DGRP-42 | 28194 | DGRP-392 | 28251 | DGRP-855 |  |  |

|  |  |  |  |  |  |
| --- | --- | --- | --- | --- | --- |
| 28129 | DGRP-59 | 28196 | DGRP-426 | 28252 | DGRP-857 |
| 28131 | DGRP-73 | 28198 | DGRP-441 | 28256 | DGRP-884 |
| 28132 | DGRP-75 | 28199 | DGRP-443 | 28257 | DGRP-890 |

**(3) RNAi screening**

For manipulation of the Gal4/UAS system, the flies were reared at 25 °C. The following strains and reagents were used for the RNAi screening:

- *da-Gal4*
- *UAS-lacZ-RNAi* (provided by Dr. R. Carthew)

The *da-Gal4* and *UAS-lacZ-RNAi* strains were backcrossed six times into the *w^1118^* genetic background before use.

**(4) UAS-RNAi strains used for screening:**

Strains obtained from VDRC are denoted with “V,” those from Bloomington with “B,” and those from NIG-Fly with “N” or “HMJ.”

The UAS-RNAi strains for candidate genes identified through the GWAS performed at 18 °C were as follows:

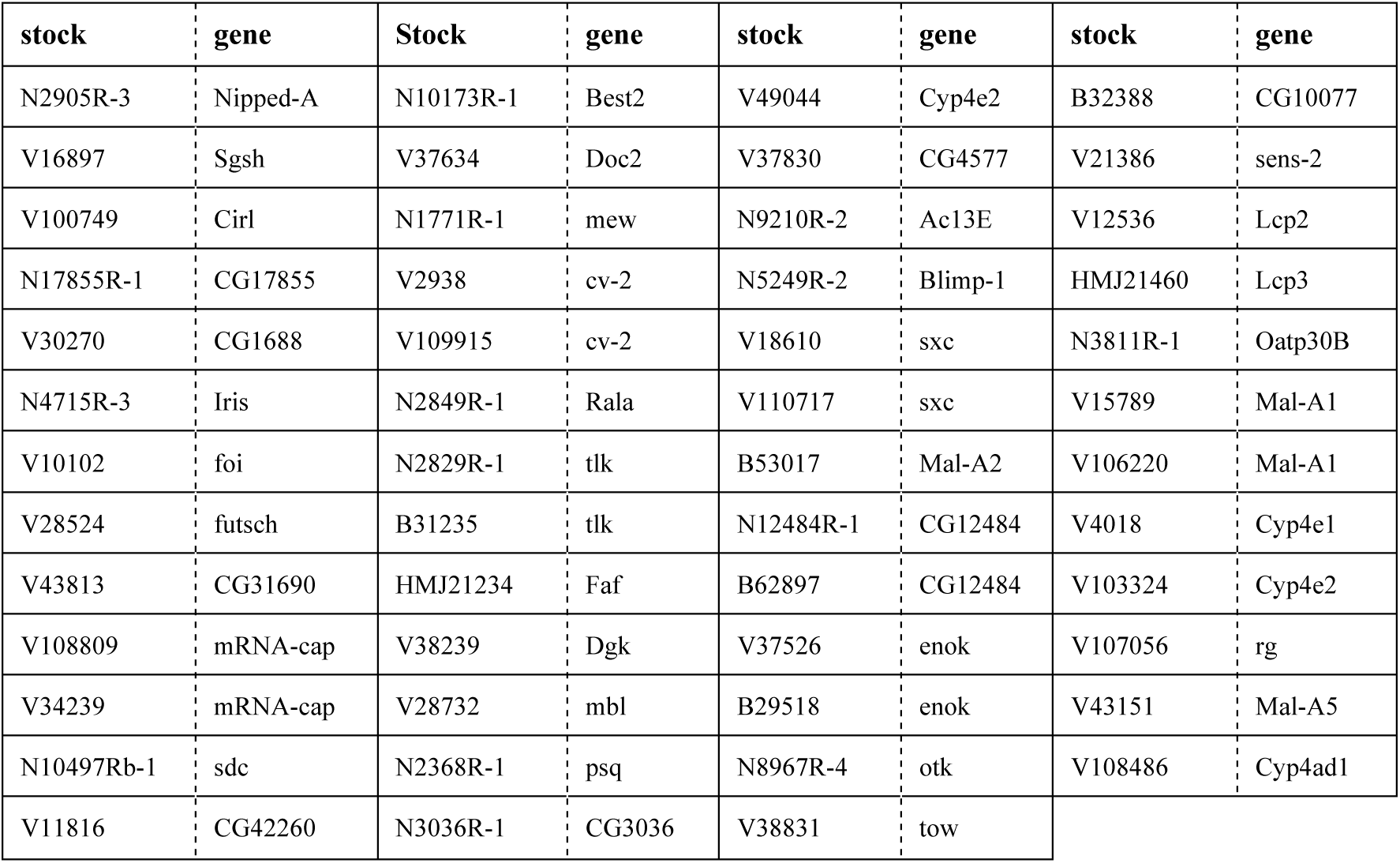

**(5) Maltase knockdown strains**

Maltase knockdown strains were used to analyze gene expression.

(5) Maltase knockdown strains Maltase knockdown strains were used to analyze gene expression.
| stock | Gene |
| --- | --- |
| V15789 | Mal-A1 |
| V106220 |  |
| V15791 | Mal-A3 |
| V7947 | Mal-A4 |

**(6) Composition of fly food**

The fly food used in this study was prepared with the following composition:

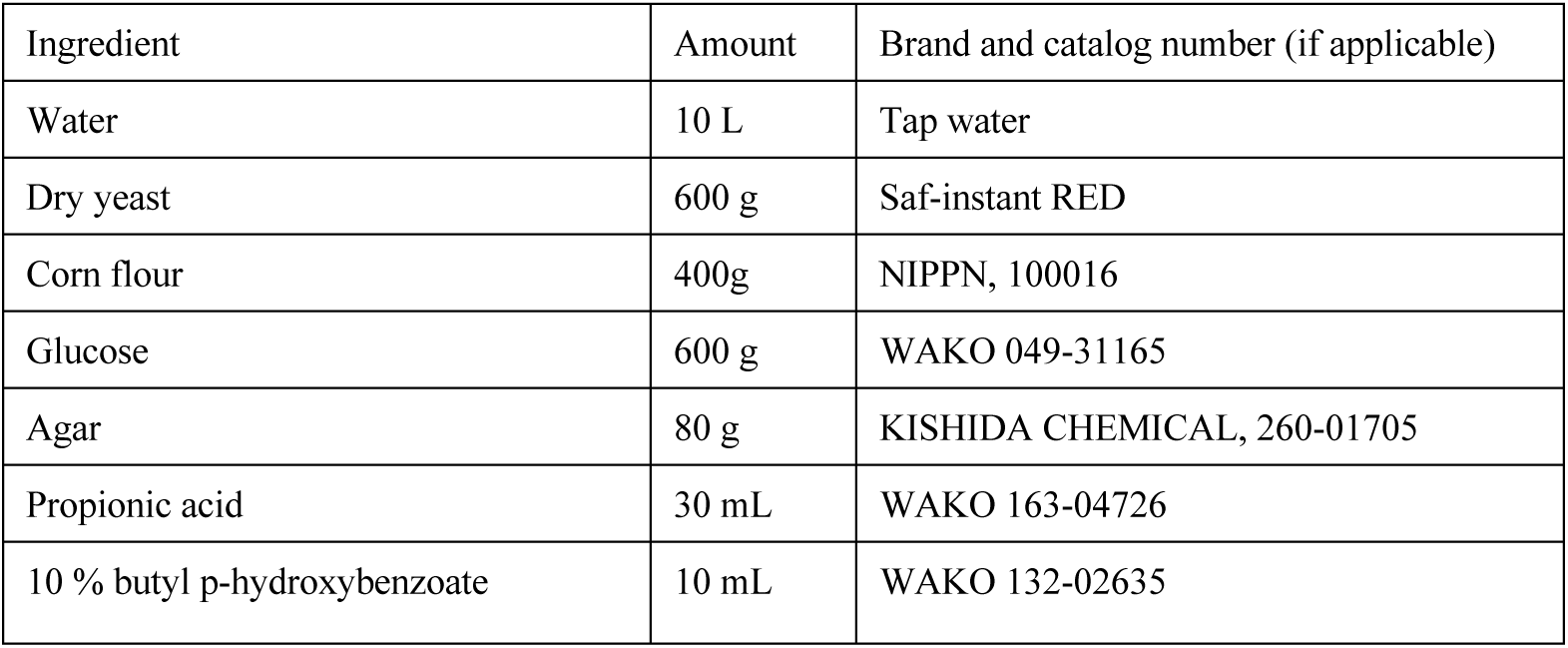

### Phenotypic analysis of DGRP lines

**(1) Counting bristle number**

Adult flies were housed in cages, and embryos younger than 24 h were collected and seeded in bottles containing approximately 30 mL of food. The flies were reared in incubators at 25 °C or 18 °C. After eclosion, the number of bristles was counted in two bottles per line. To avoid density effects on bristle counts, lines with low egg-laying rates were adjusted by increasing the number of embryos seeded in the bottles. Only the bottles containing at least 50 emerging flies were included in the analysis.

**(2) Sample collection for *Maltase* and *dilp* gene expression analysis and metabolic assays**

Embryos younger than 6 h were collected and seeded in bottles containing approximately 30 mL of food. The flies were reared in incubators at 25 °C or 18 °C. Larvae reared at 25 °C were collected at 96 h after egg laying (AEL) and those reared at 18 °C were collected at 216 h AEL. The samples were washed with 1x PBS, and three larvae were pooled per sample for whole-body analysis. For the analysis of specific tissues, the CNS was collected from 10 larvae, and 1 μL of hemolymph was extracted from 10 other larvae per sample.

**(3) Immunostaining of insulin-producing cells (IPCs) with Dilp2 antibody**

Embryos younger than 6 h were collected and seeded in bottles containing 30 mL of food. The larvae were dissected in 1x PBS at 96 h AEL (25 °C) or 216 h AEL (18 °C). The dissected samples were fixed in 4% PFA/PBS in 0.5-mL tubes, washed thrice in 0.1% PBST (0.1% Triton X-100 in PBS) for 5 min each, and blocked with PBTn (5% Normal Donkey Serum in 0.1% PBST). The samples were incubated overnight at 4 °C with the following primary antibodies diluted in PBTn: rabbit anti-Dilp2 1:500 (Li and Gong, 2015) and rabbit anti-Dilp3 1:500 (Veenstra et al., 2008). After washing four times with 0.1% PBST for 10 min each, the samples were incubated with secondary antibodies at room temperature for 2 h. After washing four times with 0.1% PBST, the samples were mounted on the SlowFade Gold Antifade Reagent (Invitrogen) for observation. IPC fluorescence was visualized using a Leica TCS-SP8 confocal microscope, and the images were processed using Fiji software. Fluorescence intensity was quantified by drawing regions of interest (ROIs) around the IPCs and calculating the mean intensities.

### Genome-wide association study (GWAS)

**(1) Data sources**

Whole-genome sequence data and genotype calls for 192 lines were obtained from the *Drosophila Genetic Reference Panel (DGRP), Freeze 2.0* (http://dgrp2.gnets.ncsu.edu/). The number of macrochaetae were assessed in females from 95 DGRP lines that thrived on our standard laboratory diet. The average number of macrochaetae per line was calculated. Information on *Wolbachia pipientis* infection status was downloaded from the DGRP website.

**(2) SNP filtering**

SNPs with call rates of < 90% or minor allele frequencies (MAF) of < 5% were excluded from the analysis. After filtering, 1,502,126 SNPs remained for the GWAS.

**(3) GWAS analysis**

An additive linear regression model was used to identify the SNPs associated with average macrochaetae number. Covariates included *Wolbachia pipientis* infection status and the first four principal components. The analysis was performed using PLINK v1.90.

### RNAi screening

For RNAi screening, five virgin *da-gal4* females were crossed with males from various UAS-RNAi lines in vials containing approximately 5 mL of food. The flies were maintained in an incubator at 25 °C. The control groups comprised *da-gal4* and UAS-RNAi flies crossed with *w^1118^* male flies. Adult female macrochaetae were counted after eclosion. To ensure consistency, UAS-RNAi lines with *sc* mutations (resulting in reduced bristle number) were compared with the *w^1118^* crossed control group. For RNAi lines from the VDRC with transgenes inserted into the Y chromosome, males with the transgene were crossed with females carrying compound X chromosomes to maintain stocks. male flies were used for bristle counting in these lines.

### Phenotypic Analysis of *Maltase* Knockdown

**(1) Bristle number count**

For knockdown analysis, five virgin *da-gal4* females were crossed with males from UAS-RNAi lines in vials containing approximately 5 mL of food and reared at 25 °C. Control groups were prepared similarly using *w^1118^* male mice. The adults were transferred to fresh vials every 5 days and maintained at 25 °C. The bristle numbers were counted after eclosion.

**(2) Sample collection for *Maltase* knockdown verification**

Three wandering larvae were collected per sample in 1x PBS from vials containing crosses of five females and three males maintained at 25 °C for 4–5 days post-crossing.

**(3) *dilp* expression and sample collection for glucose estimation**

In total, 15 females and males were housed in vials for egg-laying at 25 °C for 6 h. The larvae were collected at 96 h AEL in 1x PBS. The CNS was dissected in PBS, and 10 larval CNSs were pooled per sample. Hemolymph (2 μL) was collected from 10 larvae per sample.

### Gene expression analysis

**(1) RNA extraction**

RNA was extracted using the ReliaPrep™ RNA Tissue Miniprep System (Promega) by following the non-fibrous tissue protocol. The samples were homogenized in 250 μL LBA + TG buffer and stored at −20 °C before RNA purification.

**(2) RT-qPCR**

cDNA synthesis was performed using total RNA (500 ng for whole body or 100 ng for CNS) with the PrimeScript RT Master Mix (Perfect Real Time; Takara Bio). cDNA was diluted 10-fold, and RT-qPCR was performed using TB Green™ Premix Ex Taq™ (Tli RNaseH Plus, Takara Bio) in a QuantStudio 6 Flex Real-Time PCR system (Thermo Fisher) by following standard protocols. The PCR conditions were as follows: 45 cycles at 95 °C for 3 s and 60 °C for 30 s. PCR was followed by a melting curve analysis. The following primer sets were used for the PCR:

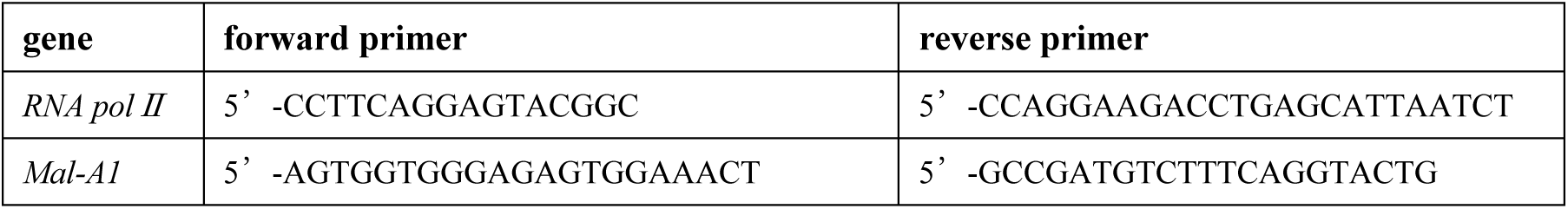
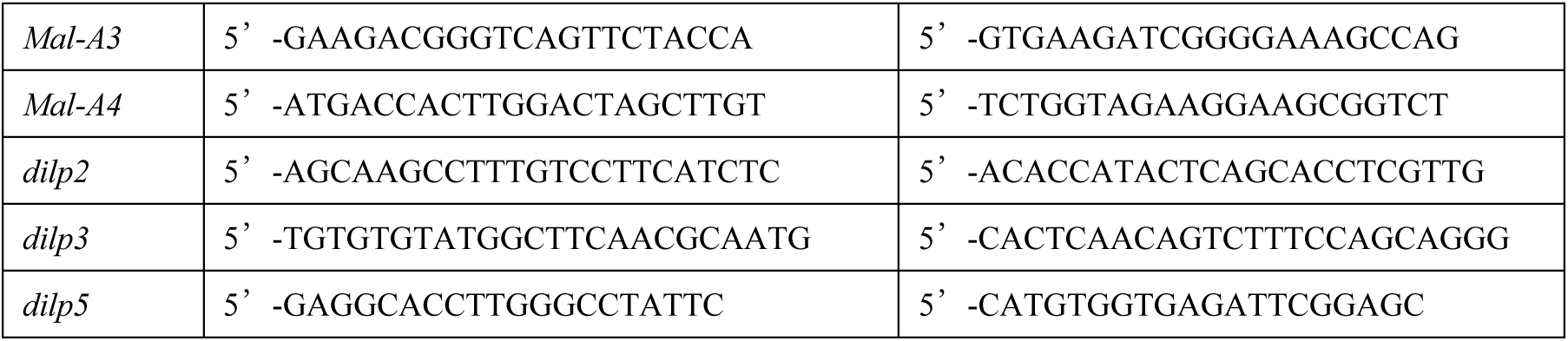

### Statistical analysis

Statistical analyses and graphical rendering were performed using the R software. Data with error bars represent mean ± standard error of the mean (SEM). For comparisons between two groups, a t-test was conducted to calculate the p-values. Statistical significance is denoted as follows: *: p < 0.05, **: p < 0.01, ***: p < 0.001, ****: p < 0.0001

### Measurement of whole-body and hemolymph metabolites using liquid chromatography–tandem mass spectrometry and gas chromatography–mass spectrometry

Widely targeted metabolomic analyses were performed as previously describe ^30^. In brief, the frozen whole-body samples (of specimens reared at 25 °C) in 1.5-mL plastic tube were homogenized for 2 min at 41.6 Hz in 300 μl of cold methanol with a single ϕ3-mm zirconia bead using a microtube homogenizer (TAITEC Corp.). Hemolymph was collected from five larvae in 1.5-ml plastic tubes, to which 300 μl of cold methanol was added. Next, the homogenates or hemolymph samples were mixed with methanol (200 μl), H2O (200 μl), and CHCl3 (200 μl) and vortexed for 20 min at room temperature. Then, the samples were centrifuged at 20,000 × *g* for 15 min at 4 °C. The insoluble pellets were used to quantify the total protein using a BCA Protein Assay Kit (Thermo Fisher Scientific). The supernatant was mixed with H2O (350 μl), vortexed for 10 min at room temperature, and centrifuged at 20,000 × *g* for 15 min at 4 °C. The aqueous phase was collected, divided into two tubes, and dried using a vacuum concentrator.

Liquid chromatography–tandem mass spectrometry (LC-MS/MS) was performed by redissolving the samples in 2 mM ammonium bicarbonate (pH 8.0). An Acquity UPLC H-Class System (Waters) was used to perform chromatographic separation under reverse-phase conditions in an ACQUITY UPLC HSS T3 column (100 mm by 2.1 mm, 1.8-μm particles, Waters) and under HILIC conditions in an ACQUITY UPLC BEH Amide column (100 mm by 2.1 mm, 1.7-μm particles, Waters). The ionized compounds were detected using a Xevo TQD triple quadrupole mass spectrometer coupled to an electrospray ionization source (Waters). The peak areas of the target metabolites were analyzed using MassLynx 4.1 software (Waters).

Gas chromatography–mass spectrometry was performed by redissolving the samples in methoxyamine pyridine solution for 90 min for oximation at 30 °C. Then, MSTFA + 1% TMCS (Thermo Fisher Scientific) was added, followed by incubation for 60 min at 37 °C for trimethylsilylation. The derivatized metabolites were analyzed using an Agilent 7890B GC coupled to a 5977A Mass Selective Detector (Agilent Technologies) in a DB-5MS + DG column (30 m × 0.25 mm, 0.25-μm film thickness; Agilent Technologies). The metabolites were detected in the selected ion monitoring (SIM) mode, and the peak areas of interest were analyzed using the QuantAnalysis software (Agilent Technologies). The metabolite signals in whole-body samples were normalized to the total protein level of the corresponding sample. The metabolite signals in the hemolymph were expressed per microliter of sample. *P* values were calculated using an unpaired two-tailed Welch’s *t* test in Microsoft Excel. Further statistical analyses such as principal component analysis were performed using MetaboAnalyst 5.0. Data were normalized to the median per sample.

## Acknowledgments

We thank the Bloomington *Drosophila* Stock Center and the Vienna *Drosophila* Resource Center for providing the fly stocks. We thank Dr. Naoya Fuse and Dr. Takashi Kamatani for discussing the GWAS. We thank Dr. Jan A Veenstra and Dr. Ryusuke Niwa for providing the Dilp3 antibody.

We thank the members of the Miura Laboratory for their technical assistance and discussions. We especially thank K. Takenaga for preparing the fly food; A. F. Kashio for the helpful comments, proofreading, and counting the bristles; and N. Shinoda and Y. Nakajima for their helpful discussions and comments.

M. M. discloses support for the research of this work from the AMED-Project for Elucidating and Controlling Mechanisms of Aging and Longevity (grant no. JP21gm5010001) and the Japan Society for the Promotion of Science (grant no. 21H04774, 23H04766, 24H00567, and 25H01842). This research was also supported by the Japan Science and Technology Agency (JST), PRESTO, under Grant Number JPMJPR24N4 to S.K., and by the joint research program of the Institute for Molecular and Cellular Regulation, Gunma University (to S.K.).

## Author contributions

S. K., I. E., and M. M. designed the experiments and wrote the manuscript. S. K. and M. M. supervised the study. S. K. and I. E. performed the experiments and analyzed the data. Y. Y. and T. N. performed metabolome analyses.

## Declaration of interests

The authors declare no competing or financial interests.

## Data availability

Data generated during the study are provided in the main article and Supplementary Information.

Additional information is available upon request from the authors.

---

**Supplementary Figure 1.**
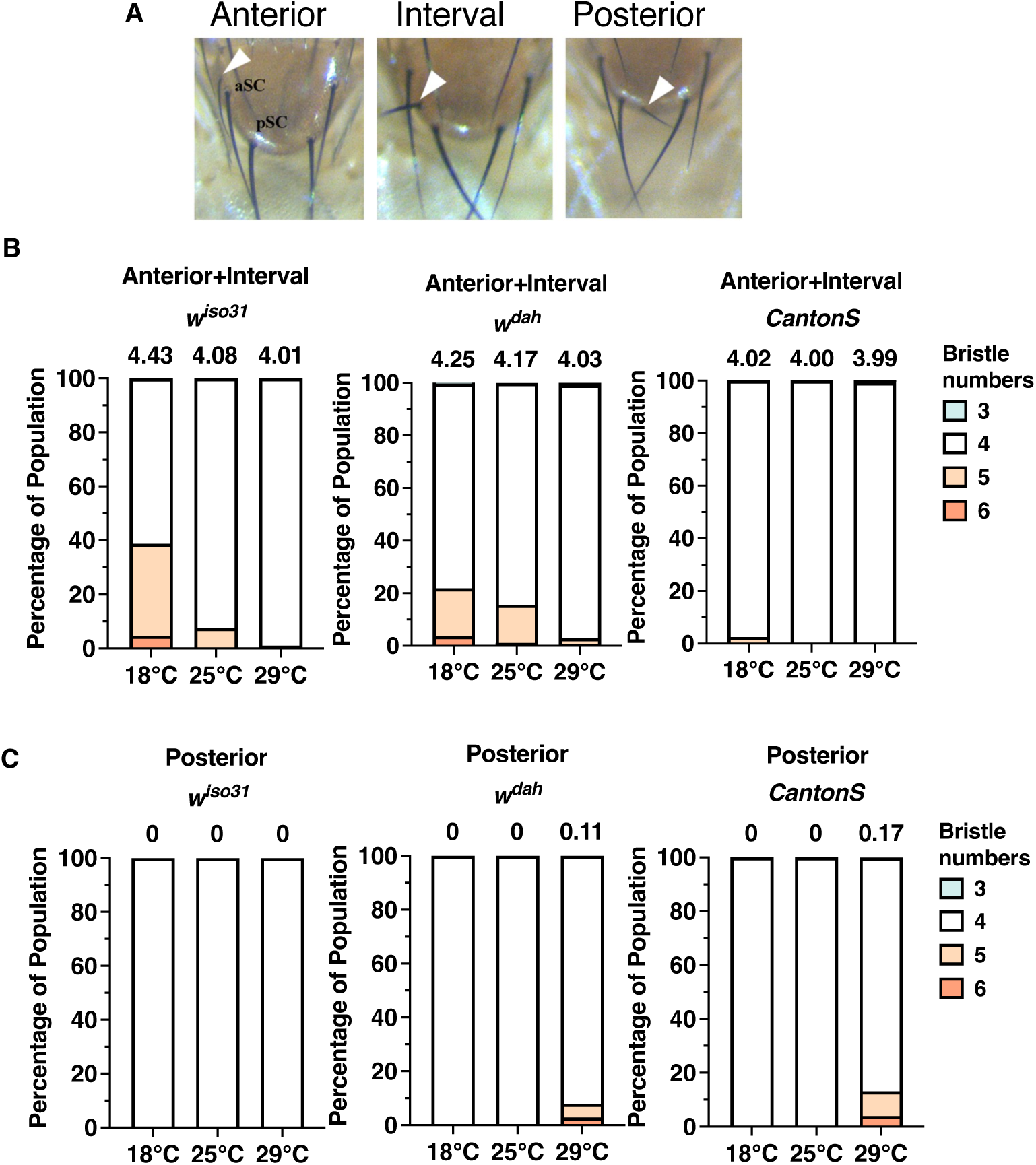
Temperature-dependent variation in location of extra bristles. **(A)** Patterns of extra bristles in *Drosophila* adult notum. These can occur around aSC (anterior), between aSC and pSC (interval), or at the lower part of pSC (posterior); (indicated as white arrowheads). **(B)** Number of bristles in females including only the extra bristles occurring in the anterior (aSC) or the interval (aSC-pSC) locations. From left to right, n=214, 2773, 95 (*w^iso31^*), 429, 198, 138 (*w^dah^*), 201, 120,152 (*Canton-S*). **(C)** Number of bristles in females with extra bristles at the lower part of pSC (posterior). The numbers above the graphs in (B) and (C) represent the mean bristle numbers and sample sizes. Data from (B) and (C) are combined in Fig. 1C. From left to right, n=214, 2773, 95 (*w^iso31^*), 429, 198, 138 (*w^dah^*), 201, 120,152 (*Canton-S*).

**Supplementary Figure 2.**
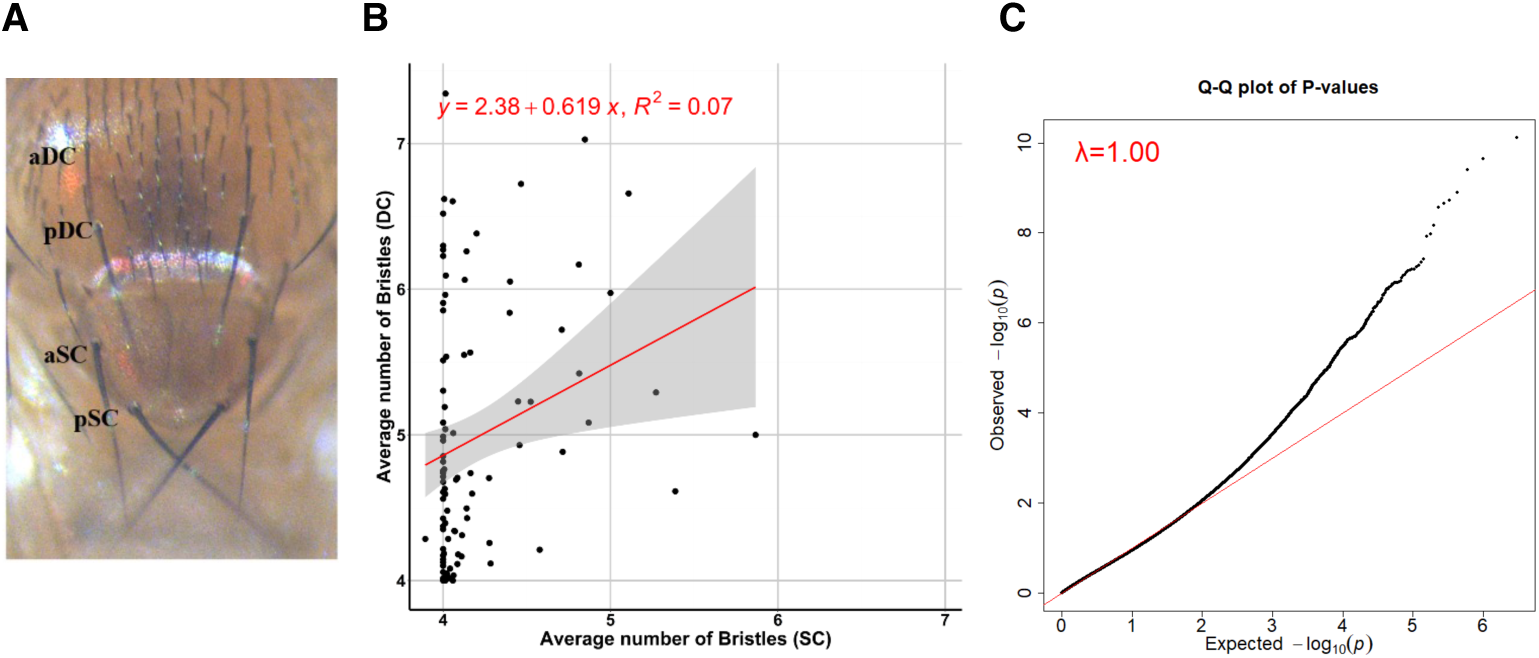
Scutella bristle numbers did not correlate with dorsal bristle numbers. **(A)** Positions of scutella bristles (aSC, pSC) and dorsal bristles (aDC, pDC). **(B)** Correlation of mean scutella brislte numbers (SC) and mean dorsal bristle numbers (DC) in DGRP strains reared at 18 °C. **(C)** Q–Q plot for mean bristle numbers at 18 °C in GWAS.

**Supplementary Figure 3.**
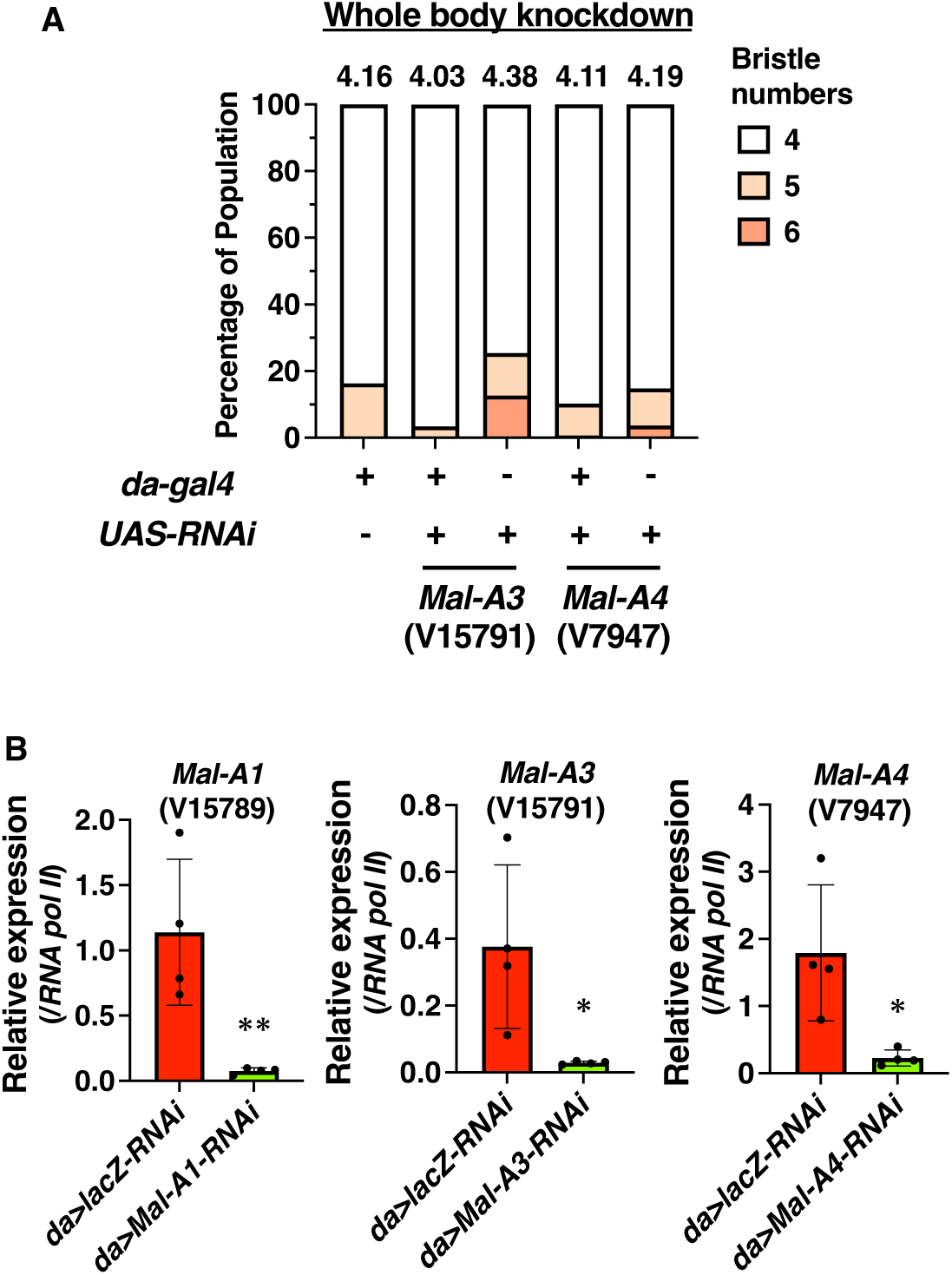
Changes in bristle numbers and gene expression following Maltase Knockdown. **(A)** Decrease in bristle numbers following whole-body knockdown of each Maltase. From left to right, n=55, 58, 55(V15791), 146, 81(V7947), 25, 55(V15794). **(B)** Knockdown efficiency of each Maltase at the whole-body level.

**Supplementary Figure 4.**
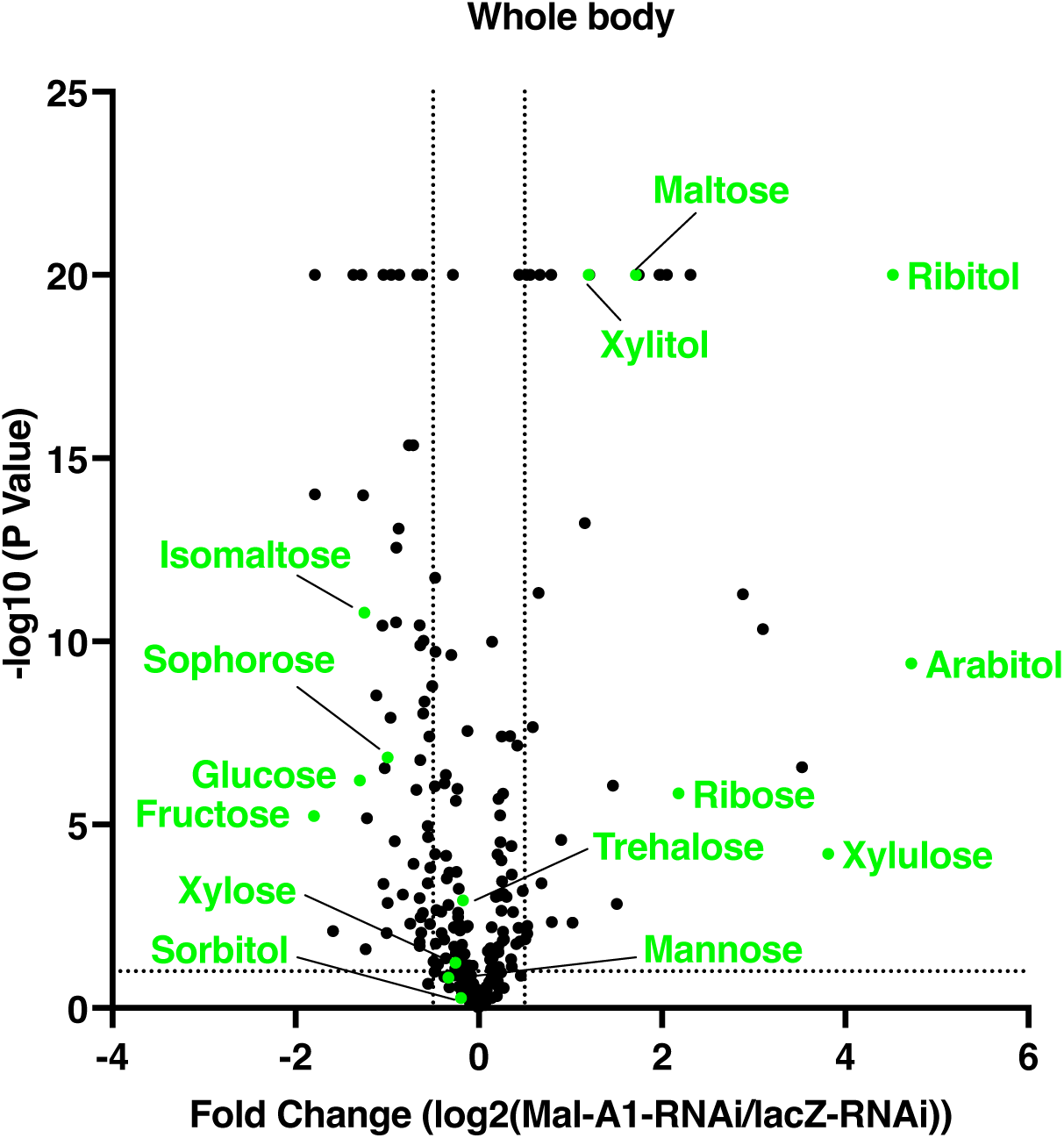
Metabolic changes induced by whole-body *Mal-A1* knockdown. Volcano plot depicting changes in metabolite abundance in whole-body samples following *Mal-A1* knockdown. Metabolites were quantified using LC–MS and GC–MS, and fold changes (*Mal-A1-RNAi* vs. *lacZ-RNAi*) are plotted against –log10(p-values). Metabolites involved in sugar metabolism are highlighted in green.

**Supplementary Figure 5.**
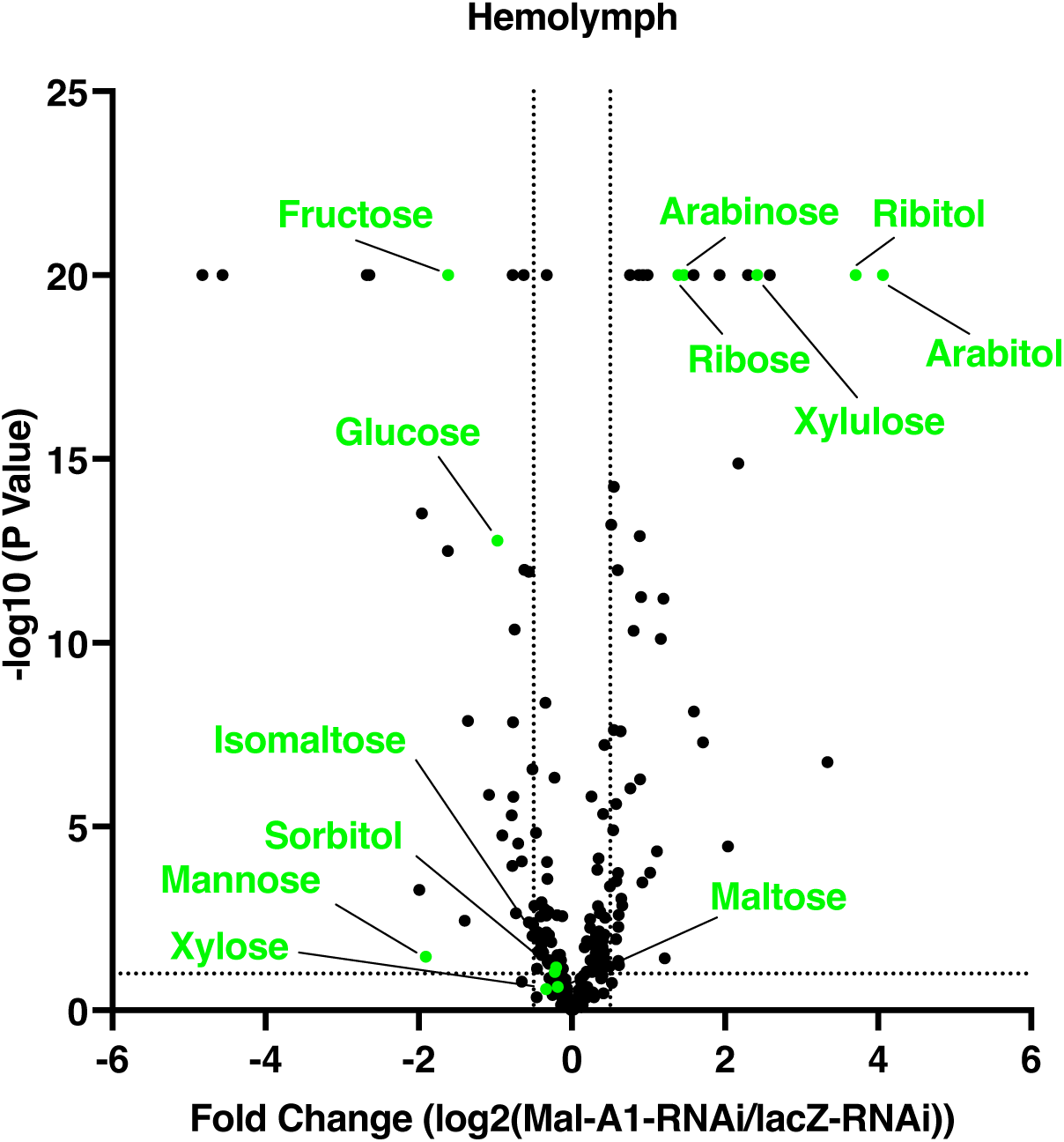
Metabolic changes induced by whole-body Mal-A1 knockdown in hemolymph. Volcano plot depicting changes in metabolite abundance in hemolymph samples following *Mal-A1* knockdown. Metabolites were quantified using LC–MS and GC–MS, and fold changes (*Mal-A1-RNAi* vs. *lacZ-RNAi*) are plotted against –log10(p-values). Metabolites involved in sugar metabolism are highlighted in green.

## Notes

### Competing Interest Statement

The authors have declared no competing interest.

